# Schemas provide a scaffold for neocortical integration at the cost of memory specificity over time

**DOI:** 10.1101/2020.10.11.335166

**Authors:** Sam Audrain, Mary Pat McAndrews

## Abstract

Memory transformation is increasingly acknowledged in theoretical accounts of systems consolidation, yet how memory quality and neural representation change over time and how schemas influence this process remains unclear. In this fMRI study, participants encoded and retrieved schema-congruent and incongruent object-scene pairs using a paradigm that probed coarse and detailed memories over 10-minutes and 72-hours. When a congruent schema was available, details were lost over time as representations were integrated in the medial prefrontal cortex (mPFC), and enhanced post-encoding coupling between the anterior hippocampus and mPFC was associated with coarser memories. Over time, pattern similarity in the hippocampus changed such that the posterior hippocampus represented specific details and the anterior hippocampus represented the general context of specific memories, irrespective of congruency. Our findings suggest schemas are used as a scaffold for accelerated consolidation of congruent information, and illustrate change in hippocampal organization of detailed contextual memory over time.

## INTRODUCTION

Memory retrieval initially depends on the hippocampus and gradually comes to be supported by the neocortex over time through the process of systems consolidation. While early theories posited that the memory trace is transferred to the neocortex regardless of episodicity (Squire and Zola-Morgan, 1991; Squire and Alvarez, 1995), there is mounting evidence that the quality of memory is related to the neurobiology that supports it. Studies of animal and human memory systems indicate that recall of detail-rich episodic memories remain dependent on the hippocampus in perpetuity, while hippocampal-neocortical dialogue promotes the strengthening of neocortical representation such that coarse, schematic memories can be supported by the neocortex independent of the hippocampus with time (Nadel and Moscovitch, 1997; Moscovitch *et al.*, 2005; Winocur, Moscovitch and Bontempi, 2010; Winocur and Moscovitch, 2011; St-Laurent *et al.*, 2014; St-Laurent, Moscovitch and McAndrews, 2016; Robin and Moscovitch, 2017; Sekeres *et al.*, 2018; Sekeres, Winocur and Moscovitch, 2018, but see Squire and Bayley, 2007; Squire *et al.*, 2015 for an alternative view). Moreover, there is evidence of specialization within the hippocampus, such that the posterior portion represents fine-grained local aspects of memory, while the anterior portion represents coarser, more global aspects (Poppenk *et al.*, 2013; Collin, Milivojevic and Doeller, 2015; Schlichting, Mumford and Preston, 2015; Robin and Moscovitch, 2017; Brunec *et al.*, 2018). Such a distinction has been proposed to arise due to intrinsic differences in hippocampal organization, as well as structural and functional connections with the neocortex along the long axis (Poppenk *et al.*, 2013; McCormick *et al.*, 2015; Adnan *et al.*, 2016; Audrain and McAndrews, 2019; Barnett, Man and McAndrews, 2019). Current theory increasingly emphasizes dynamic interactions between co-existing episodic and semantic representations of events, with the degree of detail retrieved depending on task demands and regional engagement (Winocur, Moscovitch and Bontempi, 2010; Sekeres, Winocur and Moscovitch, 2018).

Traditionally, the establishment of the neocortical memory trace was conceived of as a slow process unfolding over weeks to years, wherein the hippocampus was afforded the role of rapidly forming new orthogonal and sparsely represented memory traces, and the neocortex gradually came to extract coarser and more abstract commonalities across experiences over time (McClelland, McNaughton and O’Reilly, 1995; O’Reilly *et al.*, 2014). Indeed, memory loses precision over time in both rodents and humans, which is associated with decreased activity in the hippocampus and increased activity in the mPFC across species (Wiltgen and Silva, 2007; Winocur *et al.*, 2009; Sekeres *et al.*, 2016, 2018). This may reflect the schematization of memory over time as the memory trace becomes integrated in neocortical circuits (Nieuwenhuis and Takashima, 2011). In recent years there is increased understanding for how prior knowledge, and schemas that are extracted from multiple similar experiences in particular, can enhance memory acquisition, consolidation, and retrieval (Wang and Morris, 2010; Dudai, Karni and Born, 2015; Gilboa and Marlatte, 2017). In rodents, there is evidence that learning novel information in the context of an existing schema occurs quite quickly, effectively accelerating consolidation such that new information can be retrieved without the hippocampus faster than usual (Tse *et al.*, 2007). While schema-accelerated consolidation has yet to be definitively proven in humans, it has long been recognized that prior knowledge benefits mnemonic retention of new congruent information (Piaget, 1929; Bartlett, 1932).

Empirically, the medial prefrontal cortex (mPFC; and homologous regions in rodents) has proven to be important for schema benefits to memory across both species (Tse *et al.*, 2011; Wang, Tse and Morris, 2012; Gilboa and Marlatte, 2017; Sekeres *et al.*, 2018). There is evidence that hippocampal activity decreases (Hennies *et al.*, 2016; Sommer, 2017; Bonasia *et al.*, 2018) while mPFC activity increases during the delayed retrieval of schema-congruent relative to incongruent memories (van Kesteren *et al.*, 2010; Brod *et al.*, 2015; Sommer, 2017), indicating the increasing contribution of the mPFC in supporting schema-congruent retrieval after a period of consolidation. Further, functional coupling between the mPFC and hippocampus increases during encoding of information related to prior knowledge, and persists off-line after learning (Zeithamova, Dominick and Preston, 2012; Liu, Grady and Moscovitch, 2016, 2018; Schlichting and Preston, 2016; Sommer, 2017 c.f. van Kesteren *et al.*, 2010; Bein, Reggev and Maril, 2014), which is proposed to reflect updating of neocortical knowledge structures with related experiences (Preston and Eichenbaum, 2013; Schlichting and Preston, 2016).

Although a relative trade off in activity between the mPFC and hippocampus provides some support for the existence of schema-accelerated consolidation in humans, it is unclear exactly how this might occur. Recent theoretical accounts propose that schemas provide an organizing scaffold that new overlapping content can leverage for speeded integration (Lewis and Durrant, 2011; Gilboa and Marlatte, 2017). Such accounts rest on the idea that related memories are represented by overlapping neural ensembles in the neocortex—and the overlap of new with established content enables the rapid strengthening of new representations via Hebbian learning, presumably at the cost of memory specificity afforded by the hippocampus. The implication is that schema-congruent memories are organized and integrated according to the schema that supports acquisition, which can then be mobilized to facilitate later encoding and retrieval (Gilboa and Marlatte, 2017).

There is indeed evidence that learned arbitrary associations that share overlapping features come to be represented more similarly to each other in the mPFC than nonoverlapping events (Schlichting, Mumford and Preston, 2015; Tompary and Davachi, 2017), lending credence to the contention that overlapping information is strengthened in this region while details fall away leaving coarser, more integrated representations (Lewis and Durrant, 2011; Tompary and Davachi, 2017). Although paradigms measuring integration as a function of shared arbitrary features can speak to the role of overlap in linking episodic memories, we argue that the complex and abstracted real-world knowledge that comprise schemas likely affect representation in the brain differently. Neural population overlap as a mechanism for integration has not been examined in the context of schemas, leaving untested the idea that representational overlap accelerates consolidation. Moreover, while memory transformation and quality of memory are becoming increasingly acknowledged in theoretical accounts of long-term memory formation (Winocur, Moscovitch and Bontempi, 2010; Lewis and Durrant, 2011; Poppenk *et al.*, 2013; Preston and Eichenbaum, 2013; Dudai, Karni and Born, 2015; Sekeres, Winocur and Moscovitch, 2018; Barry and Maguire, 2019b; Yonelinas *et al.*, 2019), there is a dearth of work empirically examining how memory quality and representation change over time, and how schemas affect this process. Using resting-state fMRI, neural representational similarity analyses, and a novel behavioural paradigm sensitive to quality of memory, we targeted several lines of evidence to address the following questions: do real-world schemas act as a scaffold for the accelerated consolidation of new overlapping memories in humans, and how do the hippocampus and mPFC interact to support the consolidation and retrieval of coarse and fine-grained episodic representations in the context of schemas over time?

With these aims, participants studied a series of unique objects paired with one of four repeating scenes in an event-related fMRI paradigm (**Figure 1**), two of which were beach scenes and two of which were kitchen scenes. Half of the objects were semantically congruent with the beach or kitchen contexts (e.g. a seahorse and a beach), and half were incongruent (e.g. a pylon and a beach). Participants were subsequently shown the object cue and were asked to indicate with which schematic context the object had been paired with (beach/kitchen) in a measure of coarse retrieval, as well as with which specific scene in a measure of detailed retrieval (i.e. which beach). Memory was tested across a short delay of ten minutes and a long delay of three days, to measure change in memory quality and neural representation over time. We used a multi-voxel pattern analysis approach to quantify the degree to which overlapping schema-congruent versus incongruent memories reflect representational commonalities consistent with neural integration. We additionally collected baseline and post-encoding resting state scans to measure experience dependent changes in hippocampal-neocortical interaction as it relates to quality of subsequent memory. In this context we investigated three hypotheses that speak to the speeded integration of schema-congruent information and its organization: 1) Given that memories retrieved neocortically lack the episodic specificity of hippocampus-dependent memories, we reasoned that congruent memories should lose specificity faster than incongruent since they should be consolidated faster. 2) As post-encoding hippocampal-mPFC interaction is thought to promote updating of established memory traces, we surmised that a stronger interaction would associate with coarser memory for schema-congruent information over time. Finally, 3) we postulated that during retrieval, congruent object-scene pairs that shared the same schematic context would show greater representational overlap in the mPFC than incongruent object-scene pairs after a delay – in line with integration of congruent memories according to pre-existing schemas – but the hippocampus should maintain or differentiate distinct patterns for detailed episodic memories.

**Figure 1.**
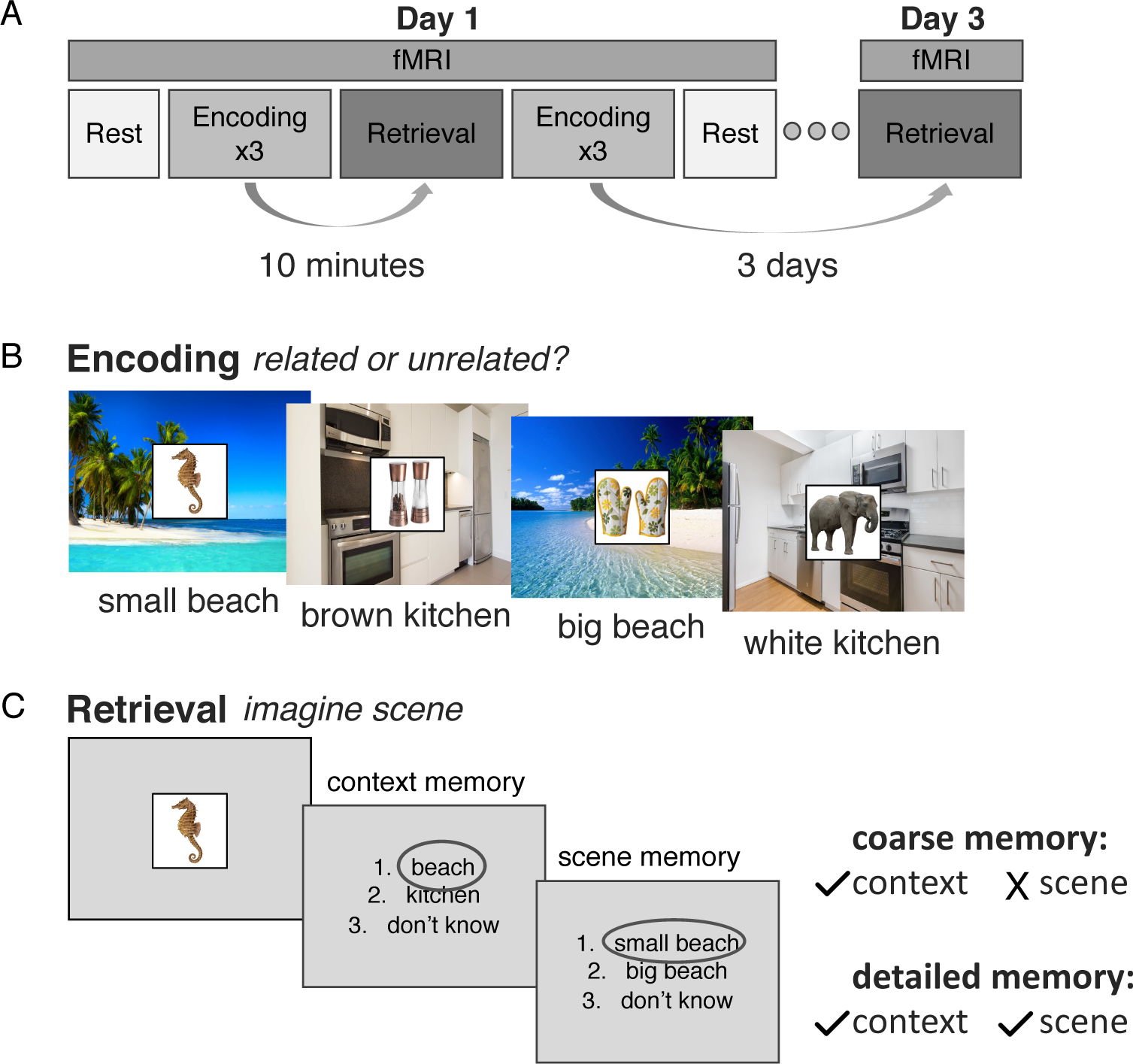
Experimental Design. A) Participants underwent encoding and retrieval sessions across a short delay of ten minutes and a long delay of three days in the fMRI scanner. Note that the delays were counterbalanced, and this figure shows only one of the task orderings. Encoding fMRI data was not analysed for the present experiment. B) During encoding participants viewed a series of object-scene pairs and indicated whether the object was related to the background scene. Participants did not view the names of the scenes during encoding but learned them prior to beginning the experiment. C) During retrieval participants were asked to imagine the scene associated with a presented object in as much detail as possible, and to indicate with which context the object had been paired with, as well as with which specific scene. Trials for which participants remembered the context a given object was paired with but not the scene were scored as coarse memories, and trials for which they remembered the context and specific scene were scored as detailed memories. Dark grey circles over responses here represent example responses.

## RESULTS

### Schema-congruent memories become coarser over time

To examine the influence of schema congruency on memory over time, memory performance was calculated for each subject as the percent of congruent or incongruent pairs in which the correct context was retrieved (regardless of the specific scene) at each delay. A linear mixed effects model indicated main effects of congruency (F(1,58)=92.41, p<0.0001) and delay (F(1,58)=83.72, p<0.0001), and a congruency x delay interaction (F(1,58)=21.61, p<0.001) on memory scores. Pairwise testing revealed that congruent pairs were remembered better than incongruent ones at the short delay (congruent: M=89.69%, SD=8.27%, incongruent M=77.82%, SD=11.16%, t(1,58)=3.99, p=0.0002). While both types of pairs where forgotten over time (congruent: t(1,58)=3.33, p=0.002, incongruent: t(1,58)=9.81, p<0.0001), congruent pairs were retained better at the long delay (congruent: M=79.82%, SD=12.16%, incongruent: M=47.39%, SD=9.91%, t(1,58)=9.91, p<0.001). The percentage of correctly remembered contexts was reliably above random chance (33.33%) for both conditions at each delay (all p<0.0001).

Next, we examined how quality of memory changed over time within each condition (**Figure 2**). At each delay, we defined detailed memories as the percent of total congruent or incongruent trials where participants correctly identified both the context (beach/kitchen) and the specific scene (big beach/small beach or brown kitchen/white kitchen) that was paired with a given object at retrieval. We defined coarse memories as the percent of total congruent and incongruent trials in which participants correctly identified the context an object had been paired with (beach/kitchen), but indicated that they did not know the specific scene, or chose the incorrect scene of the same context. We ran separate linear mixed models for coarse and detailed memories, with delay and congruency and their interaction as predictors. For detailed memories, we found a significant main effect of delay (F(1,58)=141.26, p<0.0001; short: M=66.44%, SD=17.07%; long: M=37.86%, SD=19. 57%). There was also a main effect of congruency (F(1,58)=34.00, p< 0.0001; congruent: M=60.68%, SD=21.84%, incongruent: M=46.35%, SD=28.42%). The interaction between delay and congruency for detailed memories was marginal (F(1,58)=3.56, p=0.06). For coarse memories, there were significant main effects of delay (F(1,58)=23.30, p<0.0001) and congruency (F(1,58)=14.25, p=0.0004), and a significant interaction between the two (F(1,58)=9.56, p=0.003). Pairwise testing indicated that there was no difference in the percentage of coarse congruent and incongruent memories at the short delay (congruent: M=18.18%, SD=7.74%, incongruent: 16.44%, SD=7.82%, t(1,58)=0.71, p=0.48). There were, however, more coarse congruent than incongruent memories retrieved at the long delay (congruent: M=32.24%, SD=8.93, incongruent: M=19.25%, SD=8.94%, t(1,58)=4.83, p<0.0001), which was driven by an increase in the percentage of coarse congruent memories retrieved over time (t(1,58)=5.61, p<0.0001). The percentage of coarse incongruent memories did not change over the same time period (t(1,58)=1.29, p=0.20). This pattern of results is consistent with the hypothesis that schema-congruent memories are neocortically consolidated faster than incongruent.

**Figure 2.**
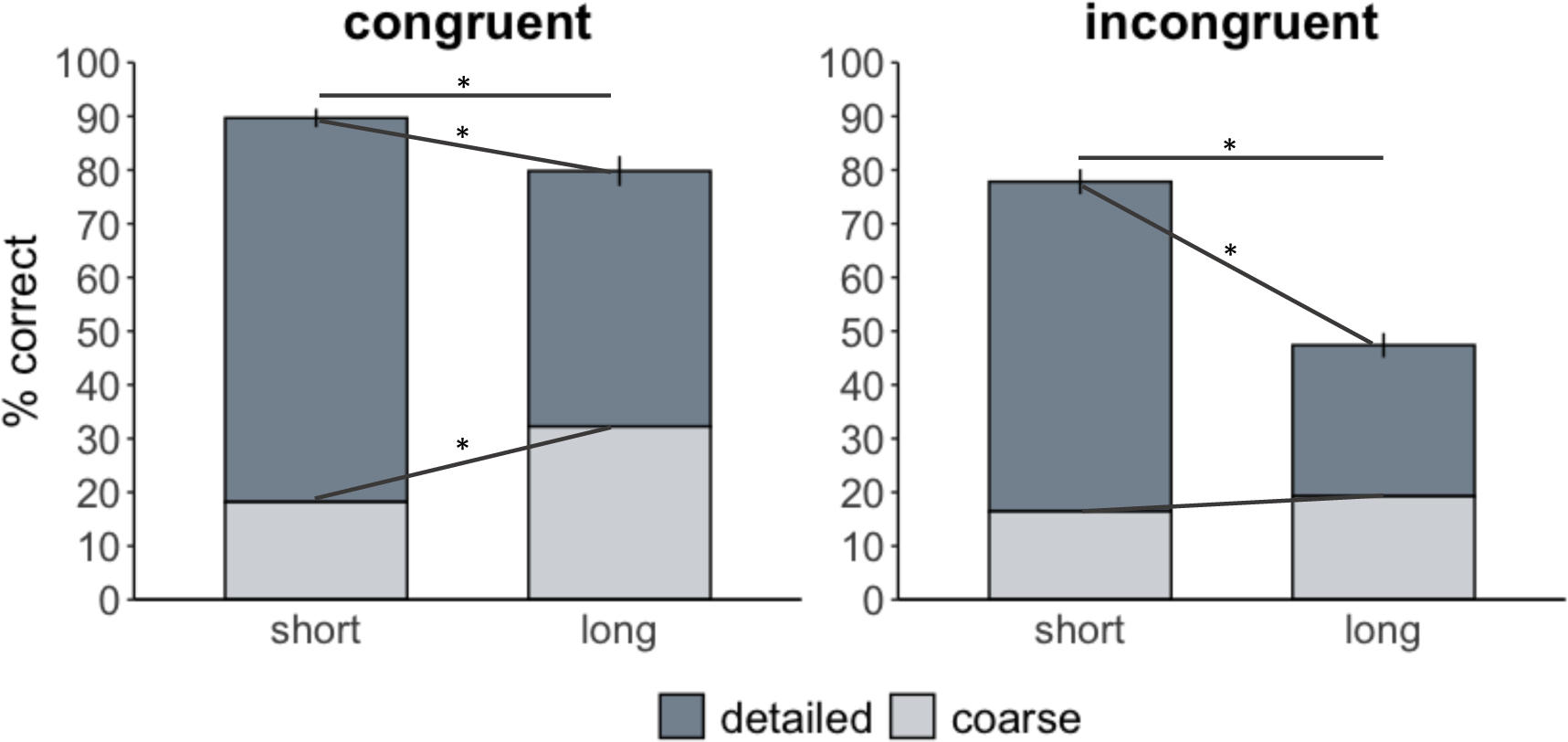
Behavioural Results. Retrieval performance as a function of congruency and memory granularity. Memories were considered coarse if participants retrieved the correct context an object had been paired with but not the specific scene and were considered detailed if they retrieved the specific scene. Error bar reflect standard error of the mean adjusted for within-subject design.*p<0.05.

As there were more detailed congruent memories retrieved than incongruent ones at the long delay, we wanted to delineate if this finding was due to a benefit for detailed memory due to schema congruency, or rather reflected the fact that there were more congruent trials retrieved to begin with. When we subtracted %retrieval at the short delay from the long delay to obtain a measure of forgetting for each subject, we found that there was no significant difference in the proportion of detailed memories forgotten between the congruent and incongruent condition (F(1,18)=2.57, p=0.13), and it was only coarse memories for which rate of forgetting differed (F(1,18)=14.47, p=0.001). Together, these behavioural findings indicate that schema-congruency benefitted memory initially and promoted better retention over time. While detailed memories were forgotten to a comparable degree in both conditions, only schema-congruent content was remembered more coarsely across a long delay.

### Anterior hippocampus – mPFC increase in post-encoding coupling is associated with coarser congruent memory over time

Prior research suggests that post-encoding connectivity in memory-relevant networks reflects early systems consolidation processes, with enhanced delayed long-term retrieval associated with increased connectivity (Tambini, Ketz and Davachi, 2010; Tambini and Davachi, 2013, 2019). In order to determine if post-encoding hippocampal-mPFC coupling was associated with coarser congruent memories over time (or in other words, with a loss of behavioural memory precision), we extracted functional connectivity between the anterior hippocampus and the mPFC at baseline and after encoding object-scene pairs for the long delay. We then subtracted each subject’s baseline from their post-encoding connectivity to acquire a measure of change in connectivity after encoding, which we correlated with behavioural memory scores across the long delay. In line with our hypothesis, we found that participants with greater post-encoding coupling between the anterior hippocampus and mPFC retrieved coarser congruent memories three days later (r=0.46, t(15)=1.98, p=0.03; **Figure 3**). Exploratory analyses indicated that connectivity between these regions did not reliably associate with detailed congruent memory across the same delay, or detailed/coarse memory for incongruent pairs (all p>0.05).

**Figure 3.**
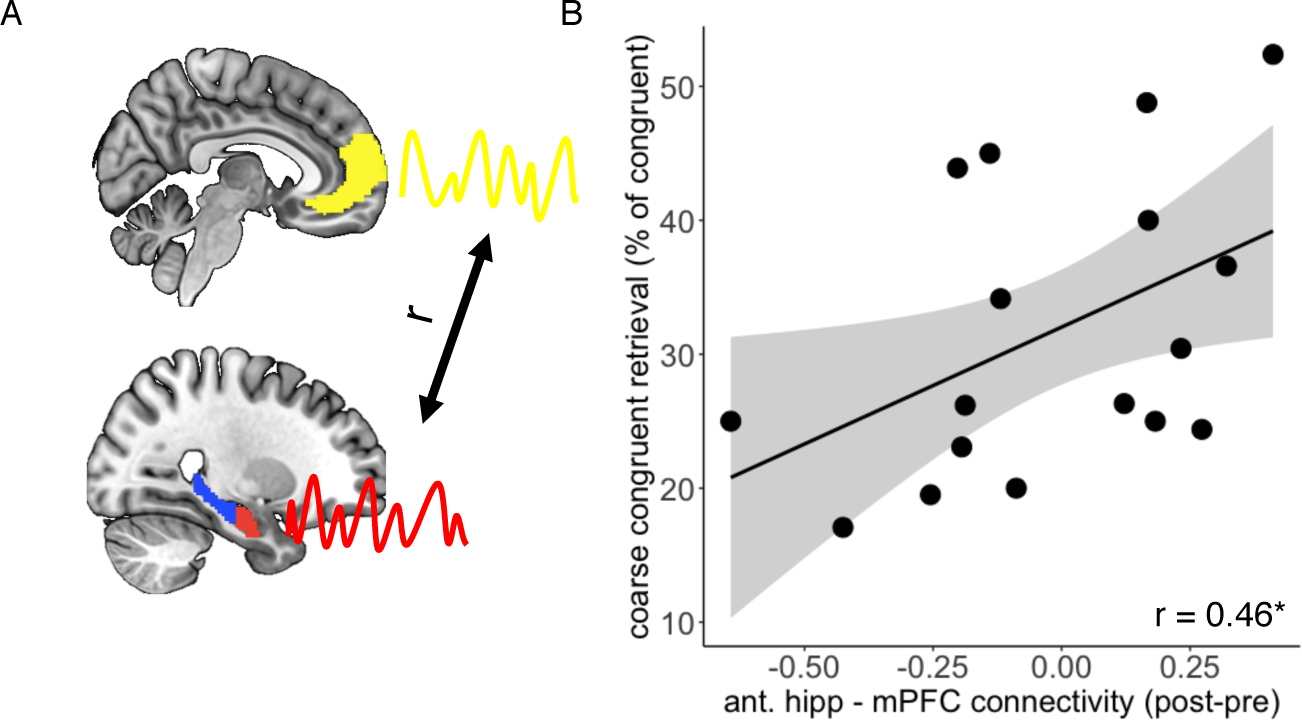
Resting state connectivity. A) Time series from the anterior hippocampus and mPFC were extracted and correlated during baseline and post-encoding rest. B) Correlation between anterior hippocampus and mPFC post-encoding coupling and coarse congruent memory scores three days layer. Coarse congruent memory was quantified as the percent of total congruent judgements for which the context was correctly retrieved but the specific scene was not. Grey ribbon represents 95% confidence interval for a one-tailed test.*p<0.05.

### Schema-congruent memories are integrated in the mPFC according to schematic context

Next we used a multi-voxel pattern analysis approach (Kriegeskorte, Mur and Bandettini, 2008; Tompary and Davachi, 2017) to quantify the degree to which schema congruent and incongruent memories share neural representations within our ROIs at each delay. We reasoned that higher pattern similarity between trials within a condition would reflect commonalities in neural representation. An increase in representational similarity over time, therefore, is consistent with the idea that there is increased neural population overlap and integration of mnemonic representations after a period of consolidation.

For congruent pairs, we extracted the pattern of voxels within the mPFC as participants were viewing a given object and successfully retrieving the associated context (regardless of memory quality), and then correlated the extracted pattern with all other patterns for congruent object-context pairs that shared the same context (**Figure 4.a**). For incongruent pairs, we computed the same set of correlations except all object-context pairs were incongruent. Despite the fact that participants are retrieving the same context (e.g. beach) in each condition, an increase in pattern similarity should be evident in the mPFC for congruent information if commonalities across object-context pairs are enhanced with consolidation due to congruency with pre-existing associations. To the extent that schemas act as a scaffold for integration, increased pattern similarity over time should be context-specific (i.e. representations for congruent object-beach pairs should become more similar to those for other congruent object-beach pairs than to patterns for congruent object-kitchen pairs). In consideration of this, we additionally correlated the pattern for each object-context pair with the patterns for all other objects that had been paired with the opposing context within each congruent/incongruent condition, so that we could decipher context specificity of neural patterns over time in a subsequent control analysis. As we were mainly interested in change in pattern similarity in the mPFC over time as it relates to congruency and did not have a priori hypotheses regarding hippocampal patterns along this dimension, we focused on the mPFC for this analysis.

**Figure 4.**
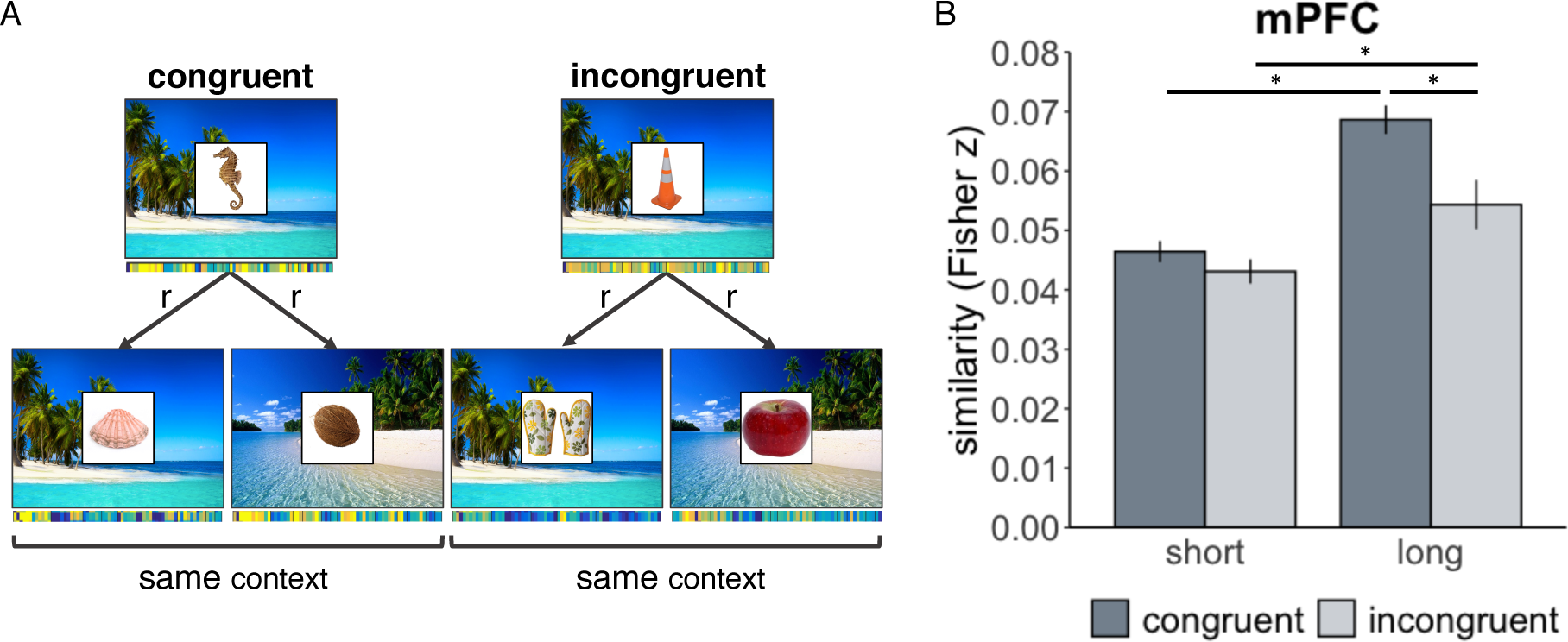
Representational similarity analysis during retrieval of congruent and incongruent object-context pairs. A) Schematic example of our analysis approach. Patterns for successfully retrieved object-context pairs (regardless of memory quality) were extracted from the mPFC and correlated within context and congruency. This example shows beach stimuli, but the same analysis was applied to kitchens. B) Resulting pattern similarity in the mPFC over time, according to congruency. Errors bar reflect standard error of the mean adjusted for within-subject design.*p<0.05. r=Pearson’s correlation.

We ran a linear mixed model predicting pattern similarity in the mPFC as a function of congruency (congruent/incongruent) and delay (short/long), with trials restricted to same-context correlations. We found main effects of congruency (t(13848)=8.74, p=0.003) and delay (t(13848)=85.18, p<0.001) as well as a significant interaction between the two (t(13848)=5.35, p=0.02; **Figure 4.b**). Pairwise testing indicated that there was no reliable difference in representational similarity of congruent and incongruent object-context pairs at the short delay (t(13848)=1.36, p=0.17). While patterns for both congruent and incongruent pairs became more similar over time (congruent: t(13848)=8.49, p<0.001; incongruent: t(13848)=2.82, p=0.005), this change was greater for the congruent pairs (long delay: t(13848)=3.51, p<0.001). In other words, object-context representations became more similar to each other in the mPFC over time if they shared a congruent context rather than an incongruent one.

We next compared these same-context correlations to across context correlations, with the hypothesis that pattern similarity would increase within context but not across context if representations were organized according to the paired schema. In other words, we expected that patterns for congruent object-beach pairs would become more similar to each other than to those for object-kitchen pairs, reflecting integration within a schematic context. We therefore computed a linear mixed model predicting pattern similarity as a function of context (same context/across context) and delay (short/long) separately for congruent and incongruent pairs. We found that for congruent pairs there was a main effect of delay, such that similarity became greater over time (F(1,17746)=108.09, p<0.001). There was also a significant main effect of context, due to greater similarity of patterns within than across context (F(1,17746)=7.16, p=0.008). There was no context by delay interaction (F(1,17746)=1.57, p=0.21). This indicates that patterns for objects that shared the same congruent context were more similar to each other than to congruent object-context pairs of the opposing context, irrespective of delay. For incongruent pairs, although there was a main effect of delay (F(1,10380)=13.98, p<0.001) whereby similarity generally increased over time, there was no effect of context (F(1,10380)=1.12, p=0.29) and no context x delay interaction (F(1,10380)<0, p=0.98). Thus, patterns for incongruent object-context pairs were becoming more similar over time but they were not being integrated within a schematic context. In fact, there was no evidence that patterns for incongruent object-beach pairs (for example) were more similar to each other than to patterns for object-kitchen pairs in the mPFC at either delay, despite the fact that these contexts were being successfully retrieved.

### The hippocampus supports representational specificity for detailed episodic memories over time

As the hippocampus is required for retrieval of specific episodic events, we next investigated whether the representation of detailed memories in the hippocampus varied as a function of the specificity and congruency of the object-scene targets. We ran a modified version of the representational similarity analysis outlined above using only items for which both the context and the specific scene were retrieved. We used the anterior and posterior hippocampus as our ROIs, given evidence that the posterior hippocampus represents more fine-grain or highly detailed information in comparison to the anterior hippocampus (Poppenk *et al.*, 2013). This time, we calculated three groups of correlations per successfully retrieved object-scene pair for each subject (**Figure 5a**). For congruent pairs, we extracted the pattern of voxels within each ROI as participants were retrieving the scene associated with a given object. We then correlated the extracted pattern with 1) the patterns of all other objects that had been paired with the same scene (and that were congruent with that scene, e.g. patterns for objects that had been paired with big beach were correlated with each other), 2) the pattern of all other objects that had been paired with the similar scene of the same context (and that were congruent with that context, e.g. patterns for objects that had been paired with big beach were correlated with patterns of objects that had been paired with small beach), and 3) the pattern of all other objects that had been paired with the other context (and that were congruent with that context, e.g. patterns for objects that had been paired with big beach were correlated with those of objects that had been paired with white kitchen and brown kitchen). We did the same thing for incongruent pairs, except all correlations were between incongruent rather than congruent trials. We submitted these correlations to a scene (same/similar/other scene) x congruency (congruent/incongruent) linear mixed model at each delay. We hypothesized that the posterior hippocampus would reflect scene specificity: patterns for objects paired with the same scene would be more similar to each other than to those for objects paired with the similar scene of the same context, as well as those for objects that had been paired with the opposite context. Conversely, we hypothesized that the anterior hippocampus would represent context but not scene specificity: correlations should be similar between representations for objects paired with the same and similar scene of the same context, but different from representations of objects that had been paired with the other context. We expected to see these patterns in the hippocampus at both delays irrespective of congruency if the hippocampus continues to represent detailed episodic content over time.

**Figure 5.**
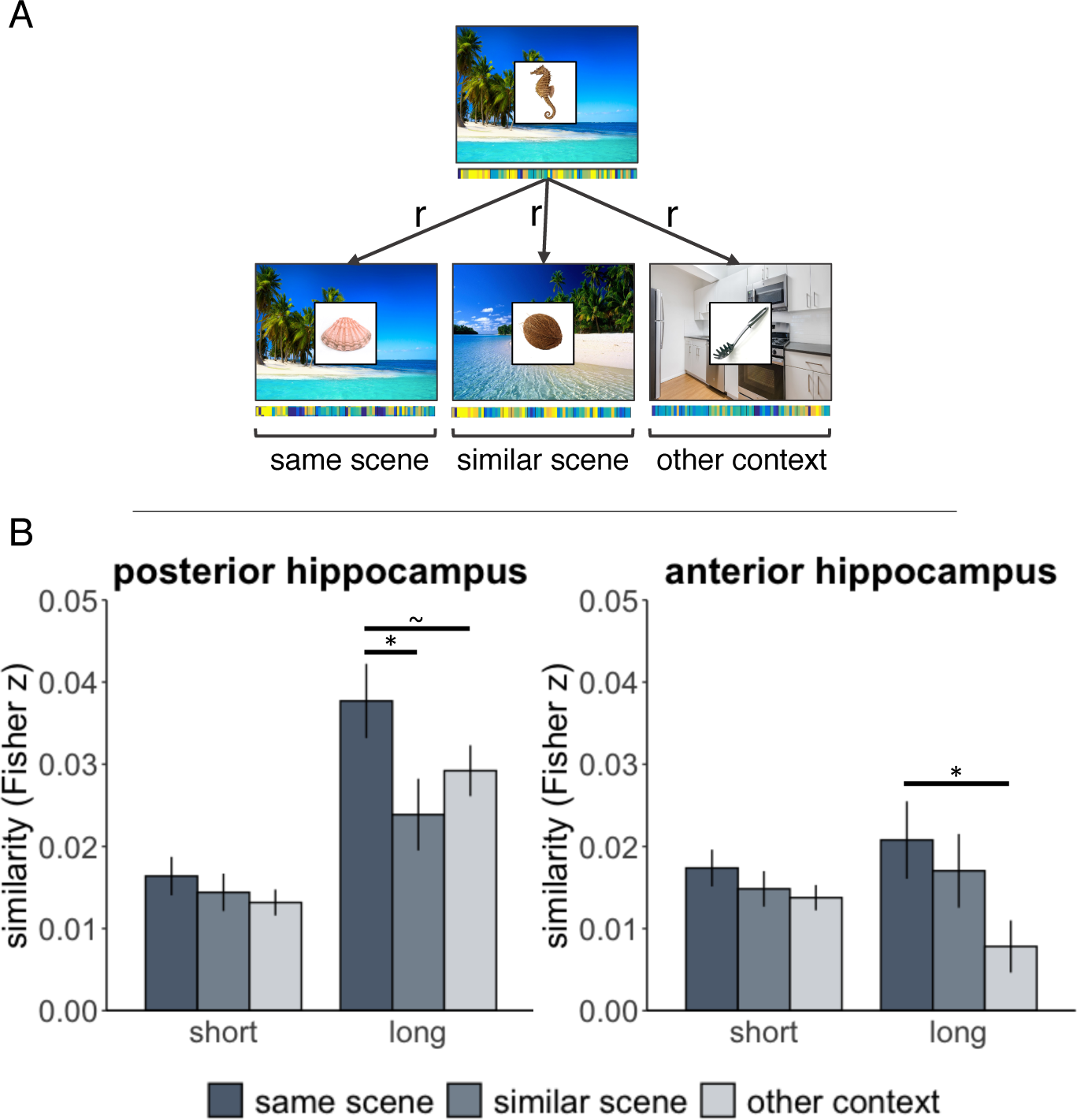
Representational similarity analysis of object-scene pairs retrieved with detail. A) Schematic example of our analysis approach. Patterns for successfully retrieved object-scene pairs were extracted from the right anterior and posterior hippocampus and correlated with objects that had shared the same scene, had been paired with the similar scene of the same context, as well as those that had been paired the other contexts (here, white kitchen is provided as an example). This example shows beach correlations but the same analysis was applied to kitchens. Note that as congruency did not interact with scene condition, we plot the main effect of scene collapsed across congruency. B) Resulting pattern similarity in the anterior and posterior hippocampus over time, according to scene/context overlap. Errors bar reflect standard error of the mean adjusted for within-subject design.*p<0.05, ∼ p=0.05.

In the posterior hippocampus there was a main effect of congruency at the short delay (F(1,12841)=8.21, p=0.004), driven by greater similarity in the incongruent than congruent condition. There was no main effect of scene (F(2,12841)=0.50, p=0.61) and no scene x congruency interaction (F(2,12841)=1.61, p=0.20), meaning that contrary to our hypothesis, patterns for objects that were paired with the same scene were no more correlated with each other than they were to objects paired with the visually similar scene or the other context scenes, regardless of congruency. At the long delay, however, a main effect of scene emerged (F(2,3284)=3.09, p=0.046). There was no main effect of congruency at the long delay (F(1,3284)=0.36, p=0.55), and no scene x congruency interaction (F(2,3284)=0.88, p=0.41), indicating that this pattern developed irrespective of congruency. As our hypothesis concerned scene granularity and there was no interaction between congruency and scene condition at either delay, we present the main effect of scene collapsed across congruency at each delay in **Figure 5b**. We further computed pairwise tests to interrogate the effect of the paired scene at the long delay (collapsed across congruency). We found greater pattern similarity between objects that shared the same scene than between objects that had been paired with similar scenes (t(3284)=2.80, p=0.005). Same scene pattern similarity was also marginally greater than pattern similarity between objects that had been paired with opposite contexts (t(3284)=1.94, p=0.05). There was no reliable difference in pattern similarity between the similar scene and other context conditions (t(3284)=1.25, p=0.21). These results indicate that when memories were retrieved with specificity the posterior hippocampus was distinguishing objects paired with the same scene from those that had been paired with other scenes over time, irrespective of congruency.

In the anterior hippocampus we also found no difference in similarity based on scene (F(2,12841)=1.12, p=0.33), congruency (F(1,12841)=1.96, p=0.16), and no scene x congruency interaction (F(2,12841)=0.13, p=0.88) at the short delay. Again, at the long delay a main effect of scene emerged (F(2,3284)=4.62, p=0.01), in the absence of an effect of congruency (F(1,3284)=0.37, p=0.55) or a scene by congruency interaction (F(2,3284)=0.15, p=0.86). As above, we plotted the effect of scene in **Figure 5b**, and tested the scene effect at the long delay with pairwise tests. While there was no difference in pattern similarity between objects that shared the same scene and those with similar scenes (t(3284)=0.56, p=0.58), or between similar and other scene correlations (t(3284)=1.56, p=0.12), there was greater similarity between patterns for objects paired with the same scene compared to that for objects paired with opposing contexts (t(3284)=2.19, p=0.03). These results indicate that over time the anterior hippocampus was not distinguishing between objects paired with specific scenes, but it was representing some degree of specificity for the overall context, regardless of congruency.

## DISCUSSION

We examined the influence of real-world schemas on systems consolidation by probing memory quality, post-encoding hippocampal-mPFC functional interaction, and representation in the mPFC and hippocampus during subsequent retrieval. We found that only schema-congruent object-scene pairs were remembered more coarsely over three days, in line with evidence for a transformation in quality of memory due to increasing reliance on neocortical retrieval over time (Winocur and Moscovitch, 2011; Sekeres, Winocur and Moscovitch, 2018), as well as better retention of (and/or greater reliance on) schematic information when detail is forgotten (Tompary, Zhou and Davachi, 2020). We further showed for the first time that the shift toward coarser quality of memory over time was associated with enhanced post-encoding coupling between the anterior hippocampus and mPFC. This finding is in line with theoretical work propounding the importance of offline functional interaction between these regions for updating established neocortical memory traces with consolidation (Preston and Eichenbaum, 2013; Schlichting and Preston, 2016; Schlichting and Preston, 2016). Finally, we present the first evidence of greater representational overlap in the mPFC during the retrieval of schema-congruent than incongruent pairs with consolidation, despite the fact that context between these two conditions was matched. Furthermore, memory representations were specifically integrated within the paired congruent schematic context, showing for the first time that schemas act as an organizing scaffold for the speeded consolidation of congruent content. As we did not find evidence of neocortical integration across these three modalities of inquiry for incongruent pairs across the same time-frame, these findings demonstrate that schemas accelerate neocortical consolidation in humans, in line with observations in rodents (Tse *et al.*, 2007, 2011; Wang and Morris, 2010; Wang, Tse and Morris, 2012).

We investigated the nature of the representations as they were influenced by delay and schema congruency, as well as their organization. Congruent and incongruent pairs shared the same overlapping contexts, but the congruent condition presumably afforded the opportunity for greater integration within a scaffold of already established object-context relationships (i.e. seashells have been experienced in the context of beaches before). Although pattern similarity in the mPFC increased over time during the retrieval of incongruent pairs that share the same context – similar to what others have reported (Tompary and Davachi, 2017) – we demonstrated a greater increase in representational overlap for congruent pairs, which was specific to paired schematic context. It follows that unique features may have been lost or minimized in both conditions over time, but it was only in the congruent condition that the learned pairs became schematically abstracted, either by strengthening overlapping elements (Lewis and Durrant, 2011; Tompary and Davachi, 2017) or by the distortion of common elements being pulled together in representational space (Milivojevic, Vicente-Grabovetsky and Doeller, 2015; Duncan and Schlichting, 2018), which is likely to occur with schematic assimilation. Alternatively, congruent memories may have become more strongly linked such that retrieval of one pair reactivated other related pairs in the neocortex (Zeithamova, Dominick and Preston, 2012), thus increasing pattern similarity across trials, although the loss of memory precision observed for congruent object-context pairs is suggestive of one of the former interpretations.

The fact that schematic context could be distinguished in the mPFC for only congruent pairs suggests that congruent object-context representations were organized according to the schema with which they were related. Thus, schematic “beach” and “kitchen” information was retrieved in the mPFC in response to congruent objects irrespective of delay. It seems as though the mPFC was not representing schematic context in the incongruent condition, despite the fact that “beach” and “kitchen” contexts were ultimately retrieved. This lack of schema context effect is at odds with the finding by Tompary & Davachi (2017) of increased representational overlap in the mPFC for arbitrary object-scene pairs within the same context relative to across contexts by one week, but it is possible that this effect emerges over longer time-frames than examined here and with slower neocortical learning (O’Reilly *et al.*, 2014). Others have observed different patterns of integration in the mPFC for arbitrary overlapping associations when such pairs were learned in a blocked manner (i.e. half of the pairs were learned first which presumably facilitated the acquisition of the subsequent trials, much like a schema would), versus when such learning proceeded in an interleaved manner as is comparable to the present experiment (Schlichting, Mumford and Preston, 2015; Schlichting and Preston, 2016). It is possible that interleaved learning (in the absence of a congruent schema) may more-so promote conjunctive coding or relational binding that facilitates inference based on specific content rather than coarse integration across trials per se (Kumaran and McClelland, 2012; Schapiro, Kustner and Turk-Browne, 2012; Zeithamova, Schlichting and Preston, 2012), which is in line with the fact that incongruent content in the present experiment did not become qualitatively coarser over time.

We also used pattern similarity analysis to examine the granularity of representations for memories retrieved with specificity in the anterior and posterior hippocampus. At the short delay objects that shared the same scene were not represented any more similarly to each other than to objects that had been paired with a different scene or context. In other words, mnemonic overlap was not mirrored with representational overlap shortly after learning the pairs. But over time, the posterior hippocampus came to represent objects that had been paired with the same scene more similarly than objects that had been paired with the different scene of the same context (e.g. big beach versus small beach), while the anterior hippocampus represented objects paired with the same scene differently from objects paired with opposing contexts (e.g. big beach vs brown kitchen or white kitchen). This finding follows rodent and recent human work indicating that by virtue of receptive field size, subfield composition, and functional and structural connectivity with the rest of the brain, the posterior hippocampus differentiates granular pieces of information in the service of episodic specificity, while the anterior hippocampus represents more global features such as episodic context (Poppenk *et al.*, 2013; Robin and Moscovitch, 2017).

It is unclear why this pattern in the hippocampus only emerged with time, but it is plausible that at the short delay visually similar overlapping information was representationally orthogonalized, in line with the well-described role of the hippocampus in pattern separation (Yassa and Stark, 2011). It was only after a period of prolonged consolidation that overlapping information came to be integrated according to the degree of contextual overlap, while also becoming differentiated from objects paired with other scenes and contexts. Notably, schematic congruency did not influence representation of these detailed memories. Several studies have reported representational changes in the hippocampus over time for overlapping or visually similar events (Milivojevic, Vicente-Grabovetsky and Doeller, 2015; Ritchey *et al.*, 2015; Favila, Chanales and Kuhl, 2016; Chanales *et al.*, 2017; Tompary and Davachi, 2017; Dandolo and Schwabe, 2018). Indeed, the most similar of these studies to ours found that pattern discriminability between overlapping and nonoverlapping object-scene pairs only emerged over time in the anterior and posterior hippocampus (Tompary and Davachi, 2017), similar to discriminability of old from new related information in the rodent hippocampus (McKenzie *et al.*, 2013). Furthermore, while a number of studies have been unable to decode mnemonic content in the hippocampus at relatively short delays (LaRocque *et al.*, 2013; Martin *et al.*, 2013; Huffman and Stark, 2014), others have documented increasing accuracy in such decoding over time (Bonnici *et al.*, 2012; Bonnici and Maguire, 2018; Lee, Kravitz and Baker, 2019), in line with the present findings. Unlike the present findings, however, some studies have decoded same versus cross context information in the hippocampus across short delays (Ritchey *et al.*, 2015; Robin, Buchsbaum and Moscovitch, 2018), especially when the contribution of different hippocampal subfields can be delineated (Kyle *et al.*, 2015; Xiao *et al.*, 2017). The finding that contextual information can be decoded in the hippocampus at the subfield level may suggest that we lack the resolution to observe such differences in the present study. Still, it is striking that representations changed so drastically over time, and it remains unclear under which conditions content can be decoded. One modern theory of memory that explicitly predicts change in representation within the hippocampus over time posits that the hippocampus reconstructs remote memories in the absence of the original trace by assembling consolidated neocortical elements into spatially coherent scenes – although we found this change occurring much sooner than these authors would have predicted (Barry and Maguire, 2019b, 2019a). Other models posit that gist predominates relative to detail over time and is mediated by the anterior hippocampus (Sekeres, Winocur and Moscovitch, 2018), but while it has been shown that the posterior hippocampus becomes less active over time (Sekeres *et al.*, 2018) it is unclear how these changes in quality of memory and activation present at the level of neural representation. Our finding that hippocampal representations change with consolidation such that objects that share the same scene share greater representational overlap while simultaneously preserving scene and context information is a novel finding in humans. Although the observed change in hippocampal representation occurred for specific memories, it is possible such memories were qualitatively different in terms of vividness or perceptual richness which was not captured by the present paradigm.

Computational modelling indicates that the hippocampus can simultaneously represent orthogonal and overlapping information (Schapiro *et al.*, 2017), in line with the proposed role of this region in both pattern separation and completion (Yassa and Stark, 2011; Rolls, 2016). Rodent work substantiates these ideas, and shows a hierarchical organization within the hippocampus that allows the simultaneous representation of related but separately acquired memories as distinct from those acquired in distinct contexts (McKenzie *et al.*, 2014). It is still unclear under which circumstances the hippocampus integrates, orthogonalizes, or separates mnemonic content, as well as the scale of such processes in humans. Studies of representational similarity in the hippocampus have generally been mixed in this regard, and have rarely been investigated in terms of change over extended delays (Duncan and Schlichting, 2018; Brunec *et al.*, 2020). Our results suggest that congruency and overlap affect representation, and the hippocampus may group common elements together while differentiating similar experiences after a period of consolidation. It follows that the direction of influence during consolidation may not be as unidirectional as once conceived (see Winocur and Moscovitch, 2011; Sekeres, Winocur and Moscovitch, 2018), and that in addition to the hippocampus driving reorganization in neocortical networks, the opposite is likely true as well – possibly through neocortical-hippocampal-neocortical loops that act to consolidate memory during sleep or awake replay of learned content (Oudiette and Paller, 2013; Klinzing, Niethard and Born, 2019; Ngo, Fell and Staresina, 2019; Rothschild, 2019), which may be especially pertinent when prior knowledge is involved (Groch *et al.*, 2017).

Finally, we did not find a loss of schema-congruent representation in the hippocampus at the long delay, as rodent work might suggest: when detailed episodic memory was retrieved, we found evidence of representational specificity in the hippocampus regardless of congruency. These observations are in keeping with trace transformation theory, which posits that even though memory quality may become less precise over time with the establishment of – and reliance on – neocortical memory traces, detailed retrieval invariably involves the hippocampus (Winocur, Moscovitch and Bontempi, 2010; Sekeres, Winocur and Moscovitch, 2018). This finding raises important considerations that have yet to be tested in the rodent literature: while schema-consistent information can be retrieved without the hippocampus relatively quickly, it is possible that such memory is nonetheless lacking the rich episodic detail and specificity the constitutes hippocampal memories (e.g. Sekeres *et al.*, 2018). Should task demands tax the memory system to retrieve such detail, we predict the hippocampus would be required – as demonstrated in rodent (and human) studies of memory in the absence of a learned schema (Winocur, Moscovitch and Bontempi, 2010; Sekeres *et al.*, 2018, 2020; Sekeres, Winocur and Moscovitch, 2018).

To conclude, we show for the first time that after encoding there is enhanced hippocampal-mPFC coupling that supports the schematization of schema-congruent memories after a prolonged period of consolidation. In parallel, we present the first evidence that real-world schemas act as organizing scaffolds that serve to accelerate the consolidation and neocortical integration of related memories. The hippocampus, on the other hand, supported specificity of representation for detailed retrieval at the long delay irrespective of congruency. Interestingly, the pattern of hippocampal representation during retrieval changed markedly over time and was suggestive of integration of overlapping content while simultaneously keeping similar memories distinct. This unexpected finding suggests that even detailed hippocampal representations change with consolidation, expanding the hypothesized role of the hippocampus to include the organization of contextual memory over time.

## Acknowledgements

We would like to thank Leanne Fernandes for her assistance with data collection, Dr. Katherine Duncan and Anuya Patil for sharing their fMRI sequence with us, Drs. Margaret Schlichting, Morris Moscovitch, and Alexander Barnett for their comments and advice during various stages of this project. This work was supported by the Natural Sciences and Engineering Research Council of Canada Research Grant #RGPIN-2015-06471 (M.P.M.) and the Toronto Neuroimaging Stimulus Grant (M.P.M. and S.A.).

## Author Contributions

Conceptualization, S.A. and M.P.M; Methodology, S.A. and M.P.M.; Software, S.A.; Formal Analysis, S.A.; Investigation, S.A.; Resources, M.P.M.; Writing – Original Draft, S.A.; Writing – Review & Editing, S.A. and M.P.M.; Visualization, S.A. and M.P.M; Supervision, M.P.M.; Funding Acquisition, S.A. and M.P.M

## METHODS

### Participants

Twenty-three young adults (8 M/15 F, mean age: 26.39 years, range: 22 – 34) participated in this experiment. Total memory score was below chance (<33%) in four participants at the long delay, and, therefore, their data for the long delay was excluded in all analyses; data at the short delay were retained. For resting state connectivity analyses an additional 2 participants were excluded as resting state scans were not collected due to technical issues. All participants were English speakers with normal or corrected to normal vision, and no active diagnosis of neurological or psychiatric disorder. The experimental protocol was approved by the University of Toronto Research Ethics Board.

### Experimental design

Participants underwent two fMRI sessions separated by approximately 72 hours. During the course of the experiment participants underwent an encoding and cued retrieval session for object-scene pairs across a 10 minute (short) delay, and again across a 72 hour (long) delay. These delays were chosen to reflect long-term memory across a relatively short and extended delay. The 72 hour delay was chosen based on behavioural piloting which indicated adequate performance across this timeframe, while also allowing multiple nights of sleep between study and test to provide the opportunity for extended consolidation processes to occur (Klinzing, Niethard and Born, 2019). All encoding and testing took place within the fMRI scanner, and the order of the delays was counterbalanced to limit confounding practice effects and differences in neural similarity that could arise due to experience with the four scenes. Participants underwent fieldmap and structural scanning during the 10 minute delay (i.e. they remained in the scanner), and went about their typical activities outside of the scanner during the 72 hour delay.

The counterbalancing procedure resulted in two groups of participants with slightly different scanning procedures. In group A, the first scanning session involved encoding object-scene pairs, along with a cued-retrieval test for the learned pairs 10 minutes later. They would then encode a new set of object-scene pairs in the scanner, to be tested 72 hours later during session 2. In group B, participants encoded object-scene pairs during the first session. During session 2, they were tested on the learned object-scene pairs (72 hour delay), and then encoded a new set of stimuli, which they were tested on 10 minutes later. All participants were administered a learning and practice test (described below) prior to scanning to prepare for the main experiment within the scanner.

#### Stimuli

Four scenic colour photos (1920 × 1080 pixels) were used in this experiment: two beaches, and two kitchens. These beaches and kitchen scenes served as backgrounds to 160 pictures of objects in white boxes (300 × 300 pixels), 60 of which were objects typically found in kitchen contexts, 60 were typically found in beach contexts, and the remaining 40 were unrelated to either context. Objects were pseudo-randomly paired with each of the four scenes within and across congruent and incongruent contexts and delay for each subject, to construct two stimulus lists per subject: one for the short and one for the long delay. Each stimulus list consisted of 80 object-scene pairs, 40 of which were congruent (20 beach objects paired with beaches, 20 kitchen objects paired with kitchens) and 40 of which were incongruent (10 beach objects paired with kitchens, 10 objects unrelated to either context paired with kitchens, 10 kitchen objects paired with beaches, 10 objects unrelated to either context paired with beaches). Half of the pairs in the incongruent condition consisted of objects typically found in the opposite context (e.g. oven mitts are typically found in kitchens, but were paired with a beach), in order to minimize the assumption that objects typically found in a context would always be paired with that context, and hence discourage the strategy of always choosing the congruent context during the memory test, described below.

#### Training Task

Given that we were interested in probing pattern similarity for scenes based on memory (and not based on re-exposure), it was important for the participants to learn the name of each of the four scenes thoroughly (“big beach”, “small beach”, “white kitchen”, “brown kitchen”) so that they could later indicate which scene was paired with an object without being visually re-presented with the scene itself during memory testing. It was equally important to ensure that participants knew the difference between scenes of the same context so that they were not inadvertently indicating the wrong scene. To that end, participants underwent a training session 1 hour prior to the first scanning session in which they learned the name of each of the four scenes, practiced visualizing the scenes in detail, and became acquainted with the task they were to undertake in the scanner. The training session consisted of 4 parts and took approximately 15 to 25 minutes to complete.

Part 1 of the training session was a self-paced format in which participants viewed each scene with its corresponding name, one at a time, and then all four scenes on the screen at once so that they could compare them. They were asked to pay attention to the name and the details of each scene, so that they would be able to name and visualize them in detail later on. In part 2 participants were shown each scene one at a time and were asked to choose the name of the scene from the four available options. They were given immediate feedback as to whether they were correct or incorrect. If they were incorrect, they repeated the process until all of the scenes were correctly named. In part 3 they were given the name of each scene one at a time, and were asked to visualize the scene in as much detail as possible, as well as to rate how vivid their visualization was on a four point scale ranging from “could not visualize” to “vivid visualization”. Right after visualization of a given scene they were asked one question pertaining to a detail of the scene (e.g. is the dishwasher located to the right or the left of the stove?), and had to select the appropriate answer out of two options. If the subject indicated they had a less than “good” visualization for any of the four scenes, or if they got any of the detail questions wrong, they re-studied the images and tried again with new questions about the details of the scenes (again, one question per scene). They repeated this process until they could produce good visualization and correctly answer the detail question for all scenes. In part 4 they underwent a practice encoding and cued-retrieval procedure as they were to be undertaken for the actual experiment for a small subset of images (12 object-scene pairs), as described in further detail below. Participants were given an abridged version of the training task before entering the scanner during the second session (72 hours later), wherein they completed Part1 and Part2 once more. This procedure served to ensure that the participants correctly remembered the name of each scene, and thus could proceed with the fMRI task.

#### Encoding

For each encoding session, participants were presented with images of 80 objects one at a time, each paired with one of four background scenes, with pairings as described above (i.e., half congruent and half incongruent). Participants’ task during encoding was to indicate if each object was related to the background scene or not. An object was to be considered “related” to the scene if the subject thought they might find the object in that context in real life. Participants were aware that they would be tested for their memory of the object-scene associations.

For each delay, they studied each object-scene pair three times across three 6.5 minute long encoding runs. All pairs were presented in each encoding run in a pseudo-random order for each subject, such that adjacent trials did not share the same scene. Participants viewed each scene for 0.1 seconds on its own before it was overlaid with the paired object for an additional 2 seconds. This brief temporal overlaying strategy was implemented to emphasize that the object and scene were separate entities rather than a unitized construct. They were then presented with a screen with response options for 1 second during which they indicated if the object-scene pair had been related or unrelated using an MRI compatible button box. The response window was followed by a jittered fixation period lasting 1, 1.5, or 2 seconds.

#### Cued retrieval

After each delay (10 minutes, 72 hours), participants underwent a cued retrieval session during which they viewed studied objects individually in the absence of the paired background scene, and were asked to retrieve the scene that had been paired with the object as vividly as possible. The 80 learned pairs were tested across four 4-minute runs with 20 objects presented in each run. Each object was presented for 2 seconds during which time participants were to visualize the paired scene. Participants were then shown a response screen and had 2 seconds to indicate with which context the object had been paired with (kitchen/beach/don’t know). The response screen remained on for the duration of the 2 seconds regardless of the speed of the button press. If they indicated that the object had been paired with a kitchen or a beach, they were then shown another response screen for an additional full 2 seconds, during which they indicated with which specific beach or kitchen scene the object had been paired with (for example, if they chose “beach” they were offered the following response options: big beach/small beach/don’t know). Piloting had revealed that some participants tended to over-rely on the “don’t know” option, so they were instructed to use this option only when they had no memory of the correct answer, in lieu of guessing (i.e. they didn’t have to have high confidence, but they should not guess). Objects were presented in a random order for each subject, and all responses were recorded using an MRI-compatible button box. Each trial ended with a jittered fixation period lasting 3 to 6 seconds.

### Behavioural data analysis

All statistical testing was performed using RStudio version 1.2.5033 (RStudio Team, 2019; http://www.rstudio.com/). To examine the influence of prior knowledge on memory over time, congruency of the object-scene pairs (congruent/incongruent) was scored based on each participant’s judgments during encoding. Given that participants viewed each object-scene pair three times during encoding, this decision was operationalized as concordance on at least two of the encoding trials. Memory performance based on congruency was calculated as the percent correct identification of the correct context (regardless of the specific scene) separately for congruent and incongruent pairs at each delay (short/long). In the examination of memory granularity, we defined detailed memories as the percent of objects for which participants correctly retrieved both the context (beach/kitchen) and the specific scene (e.g. big beach/small beach). We defined coarse memories as the percent of congruent and incongruent encoding judgements in which participants correctly identified the context with which an object had been paired with (beach/kitchen), but indicated that they did not know the specific scene, or else chose the incorrect but visually similar scene during retrieval. Differences in memory retrieval between conditions were tested using linear mixed effects models with a random intercept for each subject, using the nlme package (Pinheiro *et al.*, 2020: https://CRAN.R-project.org/package=nlme). We reported, plotted, and tested the raw descriptive means for each condition.

### fMRI parameters

All scanning was performed using a Siemens Prisma 3T full-body MRI scanner. Visual stimuli were projected onto a screen that was viewed through a mirror attached to the head coil. Functional echo-planar imaging (EPI) scans were oriented horizontally to intersect the anterior and posterior commissures (TR = 1.5s TR, TE = 26ms, flip angle = 70°, FOV = 220×220, 52 slices, 2.5mm × 2.5mm × 3mm voxels), and were acquired with a GRAPPA acceleration factor of 1, and a multiband factor of 2. Phase encoding was in the anterior to posterior direction, with interleaved acquisition in the inferior to superior direction along the z-axis. A fieldmap scan was also collected, using a double-echo gradient echo sequence with the same parameters as the EPI sequence (with the exception of the following: TR = 0.88, TE1 = 4.92ms, TE2=7.38ms, flip angle = 60°). A T1-weighted magnetization-prepared rapid-acquisition gradient echo (MPRAGE) sequence (1mm isotropic voxels, 160 sagittal slices) was also collected.

### Regions of interest definition

The right hippocampus was anatomically defined for each participant using FSL’s automatic subcortical segmentation protocol (FIRST). We chose to focus on the right hippocampus given its sensitivity to visual memory (e.g. Morris, Abrahams, Baddeley, & Polkey, 1995). Each subject’s right hippocampus was manually segmented in native space along its long axis at the uncal notch to create anterior and posterior hippocampal ROIs (Poppenk *et al.*, 2013). The mPFC mask was constructed from combining areas A14m and A10m from the Brainnettome atlas bilaterally in MNI space (https://atlas.brainnetome.org/). These ROIs are together relatively inclusive of the mPFC. We did not include some of the most ventral mPFC ROIs of the Brainnetome atlas due to a high degree of signal dropout in these areas in some of our participants, resulting in noisy signal. The resulting mPFC mask was warped into each subject’s native space using FSL’s FLIRT function.

### Resting state connectivity analysis

Pre- and post-encoding resting state scans were acquired during session 1. The baseline resting state scan was acquired at the beginning of the scan. Given that our hypotheses pertained to quality of memory over time, and given that Tompary and Davachi (2017) found that anterior hippocampus – mPFC connectivity was associated with representation of remote memories, we were specifically interested in changes in connectivity from baseline to post-encoding for stimuli that were to be tested after the long delay. The placement of the post-encoding resting-state scan occurred, therefore, directly after all three encoding runs for stimuli to be tested across the 72 hour delay (there was no resting state scan after encoding stimuli to be tested across the short delay). Rest scans were 6 minutes long, wherein participants were instructed to fixate on a small black cross in the center of a gray screen and remain awake.

Resting state scans were used to measure encoding-related changes in functional connectivity between the mPFC and long-axis hippocampal ROIs, as indexed by correlations between low frequency fluctuations in BOLD activity of each ROI (Tambini and Davachi, 2019). The resting state scans were preprocessed and modelled as separate sessions using CONN version 18b (Whitfield-Gabrieli and Nieto-Castanon, 2012; https://web.conn-toolbox.org/), which utilized the Statistical Parametric Mapping 12 (SPM12; https://www.fil.ion.ucl.ac.uk/spm/software/spm12/) toolbox via MATLAB R2016b (Mathworks) for preprocessing. The first 6 volumes were removed to allow for scanner stabilization. Motion was estimated and realignment, unwarping, and distortion correction were applied to the EPI images simultaneously. Volumes contaminated by sudden large head movements were identified using the Artifact Detection Toolbox (ART; Whitfield-Gabrieli and Nieto-Castanon, 2012), which flagged TRs with fluctuations in global signal greater than 3 standard deviations, translational motion greater than 1mm, and rotational motion greater than 0.05 radians. The EPI images were co-registered to the T1-weighted anatomical scan, and were segmented into grey matter, white matter, and cerebrospinal fluid masks for each subject. We used aCompCor (Behzadi *et al.*, 2007) to exclude physiological noise by regressing out the top five principal components from the data – as identified from a principal components analysis on the unsmoothed signal from eroded white matter and cerebral spinal fluid masks. The motion parameters (6 rigid body realignment parameters and their first order temporal derivatives, plus the high motion volumes identified by ART) were also regressed out, and the data were temporally filtered to exclude very low (<0.008 Hz) and high (>0.09 Hz) frequency fluctuations.

Average timeseries across the unsmoothed voxels within each native-space ROI were used to compute a Pearson’s correlation between the ROIs of interest for each subject (mPFC-anterior hippocampus, mPFC-posterior hippocampus). Correlation values were Fisher transformed, and the resulting values from the pre-encoding scan were subtracted from the post-encoding values for each subject. These post-pre difference scores in pairwise connectivity for each subject were then correlated with participants’ memory scores (congruent/incongruent coarse/detailed retrieval), using a one-tailed test for our a-priori hypothesis, and two-tailed tests for exploratory correlations.

### Representational similarity analysis during retrieval

#### Pattern similarity estimation

All retrieaval scans were preprocessed using FSL (FEAT; http://www.fmrib.ox.ac.uk/fsl). Encoding scans were not analyzed for the present manuscript. The first 6 volumes of the EPI images were removed to allow for scanner stabilization. For each functional run, head movement was estimated (6 rigid body motion estimates corresponding to translations and rotations around x, y, and z-axes, which were saved as regressors for later modelling) and the EPI was realigned to correct for motion. Volumes with framewise displacement > 0.9 were flagged, to be used as regressors during first level modelling in order to account for large changes in signal intensity that occur with sudden large head movements (Siegel *et al.*, 2014). To reduce spatial distortion of the EPI images, an unwrapped phase map in rad/s was constructed from the magnitude (skull-stripped) and phase fieldmap images, and applied to the EPI data simultaneously with motion correction to minimize interpolation-related image blurring. Co-registration of the EPI image to the skull-stripped T1-weighted anatomical image was also performed during this step using boundary-based registration (BBR). The EPI images were smoothed with a 3mm FWHM Gaussian kernel. All analyses took place in native space.

All preprocessed retrieval scans were modeled in each subject’s native space. We took a Least Squares Single (LSS) pattern estimation approach (Mumford *et al.*, 2012; Mumford, Davis and Poldrack, 2014), wherein each trial’s activation was estimated with a separate GLM. The first regressor in each model represented the trial of interest, and five additional regressors modeled the remaining trials within the same run according to trial type (coarse congruent, coarse incongruent, detailed congruent, detailed incongruent, forgotten). There was an additional regressor for the response window. Finally, in order to correct for head motion, there were 6 regressors for rigid body motion parameters (translations and rotations around x, y, and z-axes), as well as a regressor for each TR that was flagged as having greater framewise displacement than 0.9 during preprocessing (Siegel *et al.*, 2014). Regressors were convolved with a double gamma HRF. A map of t-values for the first parameter estimate was retained for each model and represents the activation for each trial during retrieval. For each trial, the spatial pattern of activity across each ROI was extracted into a vector and z-scored. Similarity between different vectors was calculated with Pearson correlations, which were Fisher-transformed prior to statistical testing. To avoid inflated correlations due to temporal proximity within each run, correlations were limited to trials occurring in different runs (Mumford, Davis and Poldrack, 2014).

#### RSA correlations for congruency dimension

At each delay (short/long) within-context correlations were computed on mPFC patterns between objects that shared the same context (kitchen or beach), depending on whether they were congruent or incongruent with their shared context. Specifically, similarity was computed for every retrieval trial in which the context was correctly retrieved (beach/kitchen) regardless of whether the specific scene was correctly identified (big beach/small beach or white kitchen/brown kitchen). In other words, all trials where the context was correctly remembered were included, regardless of quality of memory. We included both coarse and detailed trials in order to increase statistical power, because presumably, coarse information should be retrieved for both trial types (e.g. general features of beaches). In the congruent condition, the retrieval vector of each congruent trial (i.e. trials where the object had been congruent with its paired context) was correlated with the retrieval vectors of all other objects that shared the same context, and were also congruent with that context. Similarly, in the incongruent condition, the retrieval vector of each incongruent trial (trials where the object had been incongruent with its paired context) was correlated with the retrieval vectors of all other objects that shared the same context, and were also incongruent with that context. Given that contexts are matched in both cases, the only difference between the congruent and incongruent condition in these within-context correlations is if the objects were considered congruent or incongruent with their shared context by participants.

We additionally computed cross-context correlations to address context specificity of similarity between congruent and incongruent trials at each delay. As above, similarity was computed for every trial in which the context was correctly retrieved regardless of the specific scene. In the congruent condition, each congruent trial’s retrieval vector was correlated with the retrieval vector of all other objects that had been paired with the opposite context, and that were congruent with that context (e.g. congruent objects paired with beaches were correlated with objects that were congruent with their paired kitchens). The process was the same for the incongruent condition, in which each incongruent trial’s retrieval vector was correlated with the retrieval vector of all other objects that had been paired with the opposite context, and that were incongruent with that context. We then compared these cross-context similarity estimates to the already computed within-context similarity estimates within congruency to determine if within-context similarity was higher than cross-context in the mPFC. We additionally confirmed that differences in univariate activation between congruent and incongruent pairs over time were not driving pattern similarity results in the mPFC, as described in Supplementary Material.

#### RSA correlations for scene granularity dimension

At each delay and for each ROI (anterior hippocampus/posterior hippocampus), we computed a series of correlations between trials that had been remembered in detail, depending on congruency and the specific scene with which the object had been paired with. Retrieval similarity was computed for every retrieval trial in which both the context and the specific scene that had been paired with the object was correctly retrieved (i.e. detailed memories). For congruent object-scene pairs, each trial’s retrieval vector was correlated with 1) the retrieval vector of all other objects that had been paired with the same scene, and were congruent with that scene (**same scene correlations**), 2) the retrieval vector of all other objects that had been paired with the visually similar scene of the same context, and were congruent with that scene (**similar scene correlations**), and 3) the retrieval vector of all other objects that had been paired with the opposite context, and were congruent with that context (**other context correlations**). For the incongruent trials, we ran the same correlations except all of the correlations were between incongruent object-scene pairs (again, depending on whether the objects were paired with the same scene, visually similar scene, or other context). We additionally confirmed that differences in univariate activation between detailed congruent and incongruent pairs over time were not driving pattern similarity results in the hippocampus, as described in Supplementary Material.

#### Statistical testing of pattern similarity

Statistical testing was performed using RStudio version 1.2.5033 (RStudio Team 2019; http://www.rstudio.com/). All correlations were Fisher transformed before being submitted to statistical tests. Trial-level similarity was estimated using linear mixed effects models with a random intercept for each subject, using the nlme package (Pinheiro, Bates, DebRoy, Sarkar, & R Core Team, 2020; https://CRAN.R-project.org/package=nlme). All model assumptions were checked and verified (linearity, homogeneity of variance, normally distributed residuals). Due to differing amount of data in each condition, we reported and plotted the estimated marginal means (also known as adjusted means, extracted using the emmeans package in R: https://cran.r-project.org/web/packages/emmeans/index.html). Estimated marginal means are calculated by giving equal weight to each cell in the model, and are, therefore, unbiased by imbalances in the data; in other words, they estimate what the marginal means would be had there been equal trial numbers in each condition. Main effects and interactions were interrogated using the estimated means from each omnibus model. Within-subject error bars were computed for plotting purposes using the Morey (2008) method using the Rmisc package in R (https://cran.r-project.org/web/packages/Rmisc/index.html).

## REFERENCES

Adnan, A. et al. (2016) ‘Distinct hippocampal functional networks revealed by tractography-based parcellation’, Brain Structure and Function. Springer Berlin Heidelberg, 221(6), pp. 2999–3012. doi: 10.1007/s00429-015-1084-x.

Audrain, S. and Mcandrews, M. P. (2019) ‘Cognitive and functional correlates of accelerated long-term forgetting in temporal lobe epilepsy’, Cortex, 110, pp. 101–114. doi: 10.1016/j.cortex.2018.03.022.

Barnett, A. J., Man, V. and McAndrews, M. P. (2019) ‘Parcellation of the hippocampus using restingx functional connectivity in temporal lobe epilepsy’, Frontiers in Neurology, 10, pp. 1–12. doi: 10.3389/fneur.2019.00920.

Barry, D. N. and Maguire, E. A. (2019a) ‘Consolidating the case for transient hippocampal memory traces’, Trends in Cognitive Sciences, 23(8), pp. 635–636. doi: 10.1016/j.tics.2019.05.008.

Barry, D. N. and Maguire, E. A. (2019b) ‘Remote memory and the hippocampus: A constructive critique’, Trends in Cognitive Sciences, 23(2), pp. 128–142. doi: 10.1016/j.tics.2018.11.005.

Bartlett, F. C. (1932) Remembering: A study in experimental and social psychology. Cambridge: Cambridge University Press.

Behzadi, Y. et al. (2007) ‘A component based noise correction method (CompCor) for BOLD and perfusion based fMRI.’, NeuroImage, 37(1), pp. 90–101. doi: 10.1016/j.neuroimage.2007.04.042.

Bein, O., Reggev, N. and Maril, A. (2014) ‘Prior knowledge influences on hippocampus and medial prefrontal cortex interactions in subsequent memory’, Neuropsychologia, 64, pp. 320–330. doi: 10.1016/j.neuropsychologia.2014.09.046.

Bonasia, K. et al. (2018) ‘Prior knowledge modulates the neural substrates of encoding and retrieving naturalistic events at short and long delays’, Neurobiology of Learning and Memory, 153, pp. 26–39. doi: 10.1016/j.nlm.2018.02.017.

Bonnici, H. M. et al. (2012) ‘Detecting representations of recent and remote autobiographical memories in vmPFC and hippocampus’, Journal of Neuroscience, 32(47), pp. 16982–16991. doi: 10.1523/JNEUROSCI.2475-12.2012.

Bonnici, H. M. and Maguire, E. A. (2018) ‘Two years later – revisiting autobiographical memory representations in vmPFC and hippocampus’, Neuropsychologia, 110, pp. 159–169. doi: 10.1016/j.neuropsychologia.2017.05.014.

Brod, G. et al.. (2015) ‘Differences in the neural signature of remembering schema-congruent and schema-incongruent events’, NeuroImage, 117, pp. 358–366. doi: 10.1016/j.neuroimage.2015.05.086.

Brunec, I. K. et al.. (2018) ‘Multiple scales of representation along the hippocampal anteroposterior axis in humans’, Current Biology, 28(13), pp. 2129–2135.e6. doi: 10.1016/j.cub.2018.05.016.

Brunec, I. K. et al. (2020) ‘Integration and differentiation of hippocampal memory traces’, Neuroscience & Biobehavioral Reviews, 118, pp. 196–208. doi: 10.1016/j.neubiorev.2020.07.024.

Chanales, A. J. H. et al. (2017) ‘Overlap among spatial memories triggers repulsion of hippocampal representations’, Current Biology, 27(15), pp. 2307–2317.e5. doi: 10.1016/j.cub.2017.06.057.

Collin, S. H. P., Milivojevic, B. and Doeller, C. F. (2015) ‘Memory hierarchies map onto the hippocampal long axis in humans’, Nature Neuroscience, 18(11), pp. 1562–1564. doi: 10.1038/nn.4138.

Dandolo, L. C. and Schwabe, L. (2018) ‘Time-dependent memory transformation along the hippocampal anterior-posterior axis’, Nature Communications, 9(1), pp. 1–11. doi: 10.1038/s41467-018-03661-7.

Dudai, Y., Karni, A. and Born, J. (2015) ‘The consolidation and transformation of memory’, Neuron, 88(1), pp. 20–32. doi: 10.1016/j.neuron.2015.09.004.

Duncan, K. D. and Schlichting, M. L. (2018) ‘Hippocampal representations as a function of time, subregion, and brain state’, Neurobiology of Learning and Memory, 153, pp. 40–56. doi: 10.1016/j.nlm.2018.03.006.

Favila, S. E., Chanales, A. J. H. and Kuhl, B. A. (2016) ‘Experience-dependent hippocampal pattern differentiation prevents interference during subsequent learning’, Nature Communications, 7(1), p. 11066. doi: 10.1038/ncomms11066.

Gilboa, A. and Marlatte, H. (2017) ‘Neurobiology of schemas and schema-mediated memory’, Trends in Cognitive Sciences, 21(8), pp. 618–631. doi: 10.1016/j.tics.2017.04.013.

Groch, S. et al.. (2017) ‘Prior knowledge is essential for the beneficial effect of targeted memory reactivation during sleep’, Scientific Reports, 7(1), pp. 1–7. doi: 10.1038/srep39763.

Hennies, N. et al. (2016) ‘Sleep spindle density predicts the effect of prxior knowledge on memory consolidation’, The Journal of Neuroscience, 36(13), pp. 3799–3810. doi: 10.1523/JNEUROSCI.3162-15.2016.

Huffman, D. J. and Stark, C. E. L. (2014) ‘Multivariate pattern analysis of the human medial temporal lobe revealed representationally categorical cortex and representationally agnostic hippocampus’, Hippocampus, 24, pp. 1394–1403. doi: 10.1002/hipo.22321.

van Kesteren, Marlieke T R et al. (2010) ‘Persistent schema-dependent hippocampal-neocortical connectivity during memory encoding and postencoding rest in humans’, Proceedings of the National Academy of Sciences, 107(16), pp. 7550–7555. doi: 10.1073/pnas.0914892107.

van Kesteren, Marlieke T.R. et al. (2010) ‘Retrieval of associative information congruent with prior knowledge is related to increased medial prefrontal activity and connectivity’, Journal of Neuroscience, 30(47), pp. 15888–15894. doi: 10.1523/JNEUROSCI.2674-10.2010.

Klinzing, J. G., Niethard, N. and Born, J. (2019) ‘Mechanisms of systems memory consolidation during sleep’, Nature Neuroscience, 22, pp. 1598–1610. doi: 10.1038/s41593-019-0467-3.

Kriegeskorte, N., Mur, M. and Bandettini, P. (2008) ‘Representational similarity analysis – connecting the branches of systems neuroscience’, Frontiers in Systems Neuroscience, 2, pp. 1–28. doi: 10.3389/neuro.06.004.2008.

Kumaran, D. and McClelland, J. L. (2012) ‘Generalization through the recurrent interaction of episodic memories: A model of the hippocampal system’, Psychological Review, 119(3), pp. 573–616. doi: 10.1037/a0028681.

Kyle, C. T. et al. (2015) ‘Successful retrieval of competing spatial environments in humans involves hippocampal pattern separation mechanisms’, eLife, 4(e10499), pp. 1–19. doi: 10.7554/eLife.10499.

LaRocque, K. F. et al. (2013) ‘Global similarity and pattern separation in the human medial temporal lobe predict subsequent memory’, Journal of Neuroscience, 33(13), pp. 5466–5474. doi: 10.1523/JNEUROSCI.4293-12.2013.

Lee, S. H., Kravitz, D. J. and Baker, C. I. (2019) ‘Differential representations of rerceived and retrieved visual information in hippocampus and cortex’, Cerebral Cortex, 29(10), pp. 4452–4461. doi: 10.1093/cercor/bhy325.

Lewis, P. A. and Durrant, S. J. (2011) ‘Overlapping memory replay during sleep builds cognitive schemata’, Trends in Cognitive Sciences, 15(8), pp. 343–351. doi: 10.1016/j.tics.2011.06.004.

Liu, Z., Grady, C. and Moscovitch, M. (2017) ‘Effects of prior-knowledge on brain activation and connectivity during associative memory encoding’, Cerebral Cortex, 27, pp. 1991–2009. doi: 10.1093/cercor/bhw047.

Liu, Z., Grady, C. and Moscovitch, M. (2018) ‘The effect of prior knowledge on post-encoding brain connectivity and its relation to subsequent memory’, NeuroImage, 167, pp. 211–223. doi: 10.1016/j.neuroimage.2017.11.032.

Martin, C. B. et al.. (2013) ‘Distinct familiarity-based response patterns for faces and buildings in perirhinal and parahippocampal cortex’, Journal of Neuroscience, 33(26), pp. 10915–10923. doi: 10.1523/JNEUROSCI.0126-13.2013.

McClelland, J. L., McNaughton, B. L. and O’Reilly, R. C. (1995) ‘Why there are complementary learning systems in the hippocampus and neocortex: insights from the successes and failures of connectionist models of learning and memory.’, Psychological review, 102(3), pp. 419–457. doi: 10.1037/0033-295X.102.3.419.

McCormick, C. et al. (2015) ‘Functional and effective hippocampal-neocortical connectivity during construction and elaboration of autobiographical memory retrieval’, Cerebral Cortex, 25(5), pp. 1297–1305. doi: 10.1093/cercor/bht324.

McKenzie, S. et al. (2013) ‘Learning causes reorganization of neuronal firing patterns to represent related experiences within a hippocampal schema’, Journal of Neuroscience, 33(25), pp. 10243–10256. doi: 10.1523/JNEUROSCI.0879-13.2013.

McKenzie, S. et al.. (2014) ‘Hippocampal representation of related and opposing memories develop within distinct, hierarchically organized neural schemas’, Neuron, 83(1), pp. 202–215. doi: 10.1016/j.neuron.2014.05.019.

Milivojevic, B., Vicente-Grabovetsky, A. and Doeller, C. F. (2015) ‘Insight reconfigures hippocampal-prefrontal memories’, Current Biology, 25(7), pp. 821–830. doi: 10.1016/j.cub.2015.01.033.

Morey, R. D. (2008) ‘Confidence intervals from normalized data: a correction to cousineau (2005)’, Tutorials in Quantitative Methods for Psychology, 4(2), pp. 61–64. doi: 10.20982/tqmp.04.2.p061.

Morris, R. G. et al.. (1995) ‘Doors and people: Visual and verbal memory after unilateral temporal lobectomy.’, Neuropsychology, 9(4), pp. 464–469. doi: 10.1037/0894-4105.9.4.464.

Moscovitch et al. (2005) ‘Functional neuroanatomy of remote episodic, semantic and spatial memory: A unified account based on multiple trace theory’, Journal of Anatomy, 207(1), pp. 35–66. doi: 10.1111/j.1469-7580.2005.00421.x.

Mumford, J. A. et al. (2012) ‘Deconvolving BOLD activation in event-related designs for multivoxel pattern classification analyses’, NeuroImage, 59(3), pp. 2636–2643. doi: 10.1016/j.neuroimage.2011.08.076.

Mumford, J. A., Davis, T. and Poldrack, R. A. (2014) ‘NeuroImage The impact of study design on pattern estimation for single-trial multivariate pattern analysis’, NeuroImage. Elsevier Inc., 103, pp. 130–138. doi: 10.1016/j.neuroimage.2014.09.026.

Nadel, L. and Moscovitch, M. (1997) ‘Memory consolidation, retrograde amnesia and the hippocampal complex’, Current Opinion in Neurobiology, 7(2), pp. 217–227.

Ngo, H., Fell, J. and Staresina, B. (2019) ‘Sleep spindles mediate hippocampal-neocortical coupling during SWRs’, bioRxiv, pp. 146–151. doi: http://dx.doi.org/10.1101/712463.

Nieuwenhuis, I. L. C. and Takashima, A. (2011) ‘The role of the ventromedial prefrontal cortex in memory consolidation’, Behavioural Brain Research, 218(2), pp. 325–334. doi: 10.1016/j.bbr.2010.12.009.

O’Reilly, R. C. et al. (2014) ‘Complementary learning systems’, Cognitive Science, 38, pp. 1229–1248. doi: 10.1111/j.1551-6709.2011.01214.x.

Oudiette, D. and Paller, K. A. (2013) ‘Upgrading the sleeping brain with targeted memory reactivation’, Trends in Cognitive Sciences, 17(3), pp. 142–149. doi: https://doi.org/10.1016/j.tics.2013.01.006.

Piaget, J. (1929) The child’s conception of the world. edited by J. Tomlinson and A. Tomlinson. New York: Harcourt, Brace.

Pinheiro, J. et al. (2020) ‘nlme: Linear and Nonlinear Mixed Effects Models’. Available at: https://cran.r-project.org/package=nlme.

Poppenk, J. et al. (2013) ‘Long-axis specialization of the human hippocampus’, Trends in Cognitive Sciences, 17(5), pp. 230–240. doi: 10.1016/j.tics.2013.03.005.

Preston, A. R. and Eichenbaum, H. (2013) ‘Interplay of hippocampus and prefrontal cortex in memory’, Current Biology, 23(17), pp. R764–R773. doi: 10.1016/j.cub.2013.05.041.

Ritchey, M. et al. (2015) ‘Delay-dependent contributions of medial temporal lobe regions to episodic memory retrieval’, eLife, 4(e05025), pp. 1–19. doi: 10.7554/eLife.05025.

Robin, J., Buchsbaum, B. R. and Moscovitch, M. (2018) ‘The primacy of spatial context in the neural representation of events’, Journal of Neuroscience, 38(11), pp. 2755–2765. doi: 10.1523/JNEUROSCI.1638-17.2018.

Robin, J. and Moscovitch, M. (2017) ‘Details, gist and schema: Hippocampal–neocortical interactions underlying recent and remote episodic and spatial memory’, Current Opinion in Behavioral Sciences, 17, pp. 114–123. doi: 10.1016/j.cobeha.2017.07.016.

Rolls, E. T. (2016) ‘Pattern separation, completion, and categorisation in the hippocampus and neocortex’ Neurobiology of Learning and Memory, 129, pp. 4–28. doi: 10.1016/j.nlm.2015.07.008.

Rothschild, G. (2019) ‘The transformation of multi-sensory experiences into memories during sleep’, Neurobiology of Learning and Memory, 160, pp. 58–66. doi: 10.1016/j.nlm.2018.03.019.

RStudio Team (2019) ‘RStudio: Integrated development for R’. Boston, MA: RStudio, Inc.Available at: http://www.rstudio.com/.

Schapiro, A. C. et al. (2017) ‘Complementary learning systems within the hippocampus: A neural network modelling approach to reconciling episodic memory with statistical learning’, Philosophical Transactions of the Royal Society B: Biological Sciences, 372(1711), pp. 1–15. doi: 10.1098/rstb.2016.0049.

Schapiro, A. C., Kustner, L. V. and Turk-Browne, N. B. (2012) ‘Shaping of object representations in the human medial temporal lobe based on temporal regularities’, Current Biology, 22, pp. 1622–1627. doi: 10.1016/j.cub.2012.06.056.

Schlichting, M. L., Mumford, J. A. and Preston, A. R. (2015) ‘Learning-related representational changes reveal dissociable integration and separation signatures in the hippocampus and prefrontal cortex’, Nature Communications, 6(1), p. 8151. doi: 10.1038/ncomms9151.

Schlichting, M. L. and Preston, A. R. (2016) ‘Hippocampal–medial prefrontal circuit supports memory updating during learning and post-encoding rest’, Neurobiology of Learning and Memory, 134, pp. 91–106. doi: 10.1016/j.nlm.2015.11.005.

Schlichting, M. and Preston, A. (2016) ‘Memory integration: Neural mechanisms and implications for behavior’, Current Opinion Behavioural Science, pp. 95–121. doi: 10.1007/128.

Sekeres, M. J. et al.. (2016) ‘Recovering and preventing loss of detailed memory: differential rates of forgetting for detail types in episodic memory’, Learning & memory, 23, pp. 72–82.

Sekeres, M. J. et al.. (2018) ‘Changes in patterns of neural activity underlie a time-dependent transformation of memory in rats and humans’, Hippocampus, 28(10), pp. 745–764. doi: 10.1002/hipo.23009.

Sekeres, M. J. et al. (2020) ‘Reminders reinstate context-specificity to generalized remote memories in rats: Relation to activity in the hippocampus and aCC’, Learning & memory, 27, pp. 1–5. doi: 10.1101/lm.050161.119.

Sekeres, M. J., Winocur, G. and Moscovitch, M. (2018) ‘The hippocampus and related neocortical structures in memory transformation’, Neuroscience Letters. Elsevier, 680(August 2017), pp. 39–53. doi: 10.1016/j.neulet.2018.05.006.

Siegel, J. S. et al. (2014) ‘Statistical improvements in functional magnetic resonance imaging analyses produced by censoring high-motion data points’, Human Brain Mapping, 35, pp. 1981–1996. doi: 10.1002/hbm.22307.

Sommer, T. (2017) ‘The emergence of knowledge and how it supports the memory for novel related information’, Cerebral Cortex, 27, pp. 1906–1921. doi: 10.1093/cercor/bhw031.

Squire, L. R. et al. (2015) ‘Memory consolidation’, Cold Spring Harbor Perspectives in Biology, 7(8), pp. 1–22. doi: 10.1101/cshperspect.a021766.

Squire, L. R. and Alvarez, P. (1995) ‘Retrograde amnesia and memory consolidation: A neurobiological perspective’, Current Opinion in Neurobiology, 5, pp. 169–177.

Squire, L. R. and Bayley, P. J. (2007) ‘The neuroscience of remote memory’, Current Opinion in Neurobiology, 17, pp. 185–196. doi: 10.1016/j.conb.2007.02.006.

Squire, L. R. and Zola-Morgan, S. (1991) ‘The medial temporal lobe memory system’, Science, 253(5026), pp. 1380–1386.

St-Laurent, M. et al. (2014) ‘The perceptual richness of complex memory episodes is compromised by medial temporal lobe damage’, Hippocampus, 24, pp. 560–576. doi: 10.1002/hipo.22249.

St-Laurent, M., Moscovitch, M. and McAndrews, M. P. (2016) ‘The retrieval of perceptual memory details depends on right hippocampal integrity and activation’, Cortex, 84, pp. 15–33. doi: 10.1016/j.cortex.2016.08.010.

Tambini, A. and Davachi, L. (2013) ‘Persistence of hippocampal multivoxel patterns into postencoding rest is related to memory’, 110(48). doi: 10.1073/pnas.1308499110.

Tambini, A. and Davachi, L. (2019) ‘Awake reactivation of prior experiences consolidates memories and biases cognition’, Trends in Cognitive Sciences, 23(10), pp. 876–890. doi: 10.1016/j.tics.2019.07.008.

Tambini, A., Ketz, N. and Davachi, L. (2010) ‘Enhanced brain correlations during rest are related to memory for recent experiences’, Neuron, 65, pp. 280–290. doi: 10.1016/j.neuron.2010.01.001.

Tompary, A. and Davachi, L. (2017) ‘Consolidation promotes the emergence of representational overlap in the hippocampus and medial prefrontal cortex’, Neuron. Elsevier Inc., 96(1), pp. 228–241.e5. doi: 10.1016/j.neuron.2017.09.005.

Tompary, A., Zhou, W. and Davachi, L. (2020) ‘Schematic memories develop quickly, but are not expressed unless necessary’, PsyArXiv. doi: 10.31234/osf.io/k4fea.

Tse, D. et al. (2007) ‘Schemas and memory consolidation’, Science, 316(5821), pp. 76–82. doi: 10.1126/science.1135935.

Tse, D. et al.. (2011) ‘Schema-dependent gene activation and memory encoding in neocortex’, Science, 333(6044), pp. 891–895. doi: 10.1126/science.1205274.

Wang, S. and Morris, R. G. M. (2010) ‘Hippocampal-neocortical interactions in memory formation, consolidation, and reconsolidation’, Annual Review of Psychology, 61, pp. 49–79. doi: 10.1146/annurev.psych.093008.100523.

Wang, S., Tse, D. and Morris, R. G. M. (2012) ‘Anterior cingulate cortex in schema assimilation and expression’, Learning & memory, 19, pp. 315–318. doi: 10.1101/lm.026336.112.

Whitfield-Gabrieli, S. and Nieto-Castanon, A. (2012) ‘Conn: a functional connectivity toolbox for correlated and anticorrelated brain networks.’, Brain connectivity, 2(3), pp. 125–41. doi: 10.1089/brain.2012.0073.

Wiltgen, B. J. and Silva, A. J. (2007) ‘Memory for context becomes less specific with time’, Learning and Memory, 14(4), pp. 313–317. doi: 10.1101/lm.430907.

Winocur, G. et al. (2009) ‘Changes in context-specificity during memory reconsolidation: selective effects of hippocampal lesions’, Learning and Memory, 16(11), pp. 722–729. doi: 10.1101/lm.1447209.

Winocur, G. and Moscovitch, M. (2011) ‘Memory transformation and systems consolidation’, Journal of the International Neuropsychological Society, 17(05), pp. 766–780. doi: 10.1017/S1355617711000683.

Winocur, G., Moscovitch, M. and Bontempi, B. (2010) ‘Memory formation and long-term retention in humans and animals: Convergence towards a transformation account of hippocampal-neocortical interactions’, Neuropsychologia, 48(8), pp. 2339–2356. doi: 10.1016/j.neuropsychologia.2010.04.016.

Xiao, X. et al.. (2017) ‘Transformed neural pattern reinstatement during episodic memory retrieval’, Journal of Neuroscience, 37(11), pp. 2986–2998. doi: 10.1523/JNEUROSCI.2324-16.2017.

Yassa, M. A. and Stark, C. EL (2011) ‘Pattern separation in the hippocampus’, Trends in Neurosciences, 34(10), pp. 515–525. doi: 10.1038/jid.2014.371.

Yonelinas, A. P. et al.. (2019) ‘A contextual binding theory of episodic memory: Systems consolidation reconsidered’, Nature Reviews Neuroscience, 20, pp. 364–375. doi: 10.1038/s41583-019-0150-4.

Zeithamova, D., Dominick, A. L. and Preston, A. R. (2012) ‘Hippocampal and ventral medial prefrontal activation during retrieval-mediated learning supports novel inference’, Neuron, 75(1), pp. 168–179. doi: 10.1016/j.neuron.2012.05.010.

Zeithamova, D., Schlichting, M. L. and Preston, A. R. (2012) ‘The hippocampus and inferential reasoning: Building memories to navigate future decisions’, Frontiers in Human Neuroscience, 6, pp. 1–14. doi: 10.3389/fnhum.2012.00070.

