## Supplementary Material for "Schemas provide a scaffold for neocortical integration at the cost of memory specificity over time"

### Univariate control analyses

To exclude the possibility that differences in univariate activation across congruency and delay were driving pattern similarity results (Dimsdale-Zucker & Ranganath, 2018), we extracted the univariate activation averaged across ROI (mPFC/anterior hippocampus/posterior hippocampus) for each trial, and modelled univariate estimates as a function of congruency of the trials (congruent/incongruent) and delay (short/long) separately for each ROI. To be consistent with the analyses in the main manuscript, we included the beta estimates for all correctly retrieved trials regardless of quality of memory in our mPFC model, and all correctly retrieved detailed memory trials in our anterior and posterior hippocampus models. Should any effects on univariate activation mirror the effects observed in our RSA analyses it might indicate that such effects were driven by differences in univariate activation rather than pattern similarity. In the mPFC there was no effect of delay ($F(1,2435) = 0.34$, $p = 0.56$), condition ($F(1,2435) = 2.20$, $p = 0.14$), or an interaction between the two ($F(1,2435) = 2.06$, $p = 0.14$) on trial-wise activation. Likewise, univariate activation for trials retrieved with detail did not reliably vary as a function of congruency or delay in the anterior (congruency: $F(1,1737) = 0.04$, $p = 0.84$; delay: $F(1,1737) = 0.05$, $p = 0.83$; interaction: $F(1,1737) = 0.22$, $p = 0.64$) or posterior hippocampus (congruency: $F(1,1737) = 1.71$, $p = 0.19$; delay: $F(1,1737) = 0.49$, $p = 0.49$; interaction: $F(1,1737) = 0.43$, $p = 0.51$). It is therefore unlikely that any of our results are driven by differences in univariate activation based on congruency of the object-scene pairs or delay.
